# Alterations in swimming behavior of *Daphnia* exposed to polymer and mineral particles: towards understanding effects of microplastics on planktonic filtrators

**DOI:** 10.1101/406587

**Authors:** E. Gorokhova, O. Könnecke, M. Ogonowski, Z. Gerdes, AK Eriksson Wiklund

## Abstract

Concerns have been raised that microplastics (MP) can impact aquatic organisms by compromising their nutrition. However, little is understood about the mechanisms of the adverse effects of MP in suspension-feeders that routinely ingest particles of low nutritional value, such as mineral particles. We compared effects of non-edible particles (MP and kaolin) mixed with microalgae on the swimming and filtering behavior of a planktonic filtrator *Daphnia magna*; incubations with only algae served as controls. The following questions were addressed: (1) Are there differences in swimming movements between the daphnids exposed to MP and those exposed to kaolin? and (2) Whether occurrence of biofilm on the particle surface affects daphnid swimming and how these effects differ between the kaolin- and MP-exposed animals? We found that both kaolin and MP altered swimming, yet in opposite way, with a decrease of filtration-related movements in kaolin and their increase in MP. The difference was amplified in biofilm coated particles, indicating that daphnids spend more energy when swimming in suspension with MP, and even more when the MP have biofilm. The increased swimming activity of filtrators exposed to plastic litter decaying to microparticles may translate into changes in energy balance and growth.

## 1. Introduction

Microplastics (MP) have recently been recognized as emerging environmental contaminants. However, the potential impacts of MP for aquatic animals are not well understood, although they are commonly referred as causing various feeding disturbances. A major methodological issue related to testing MP effects is the inadequacy of standard ecotoxicological methods that have low sensitivity and little relevance for detecting ecologically likely effects (Ogonowski et al. 2016).

In nature, filter- and suspension-feeders are the most likely consumers capable of ingesting MPs. These animals are also an important “entrance” for MP to aquatic food webs because they are at the lower trophic level, where the ingestion of MPs is most probable. Experimental studies have shown that filter- and suspension-feeders frequently ingest different types of MPs, with concomitant effects on food intake, growth, and reproduction. These effects, however, result from the food dilution with nutritiously inert material and do not as such represent a toxic response. Moreover, in such experiments, commercially available plastic beads, so-called virgin MPs, are commonly used. Such particles have no coating with dissolved organic material or microorganisms (biofilms), which may affect interactions between the particle and filtering apparatus. When MP or any other particle are exposed to natural plankton assemblages or sediment communities, their surfaces become coated with various organic substances, such as carbohydrates or peptides, and colonized by various microorganisms, such as bacteria, algae, fungi and protists, building complex biofilms. Therefore, it is relevant to understand how filtrators interact with biofilm-covered MP and whether these interactions are different from those with virgin MP commonly used in the experiments.

Motion analysis is a set of techniques to quantify movement patterns and behavioral responses to external stimuli. The application of motion analysis in aquatic ecotoxicology addresses effects of abiotic stressors and incorporates data quantifying movements at a range of frequencies, thus providing a holistic analysis of animal motion. This type of analytical approach is applicable to a wide range of species, at various developmental stages. Some types of movements can be particularly informative for a given organism, but might also be an advantage to incorporate a range of movements into a single analysis (Tills et al. 2013). This approach is particularly useful for analyzing movements of a filter-feeder exposed to edible and non-edible particles and evaluating whether MPs can be perceived differently from other naturally occurring particles, such as clay, lignin, and cellulose, and can affect filtration behavior in other ways than, for example, mineral particles.

We studied swimming behavior of the cladoceran *Daphnia magna*, a common non-selective filter-feeder and a model species in ecology and ecotoxicology. The daphnids were exposed to the natural clay (kaolin) and MP of similar size. Moreover, both particle types were coated with protein to produce an artificial biofilm that would change surface properties and make our experimental particles more relevant to the naturally occurring particles. To characterize surface properties, we measured zeta potential, which is an important parameter affecting particle retention on the feeding appendages of various filter feeders (Matz and Jurgens 2001). The particles are only captured by a feeding apparatus of a filter-feeder when the Reynolds number is low due to the laminar flow and slow fluid movement (Gerritsen and Porter 1982). This implies that particles having different zeta potential, due to the nature of the material and/or occurrence of the biofilm, will have different retention capacity and thus the probability of being ingested. Therefore, the surface charge needs to be taken into account when studying interactions between MP and filtering mechanisms.

The following questions were addressed:

1. Are there differences in swimming movements between the daphnids exposed to MP and those exposed to kaolin;
2. Whether occurrence of biofilm on the particle surface affects zeta potential and daphnid swimming; and
3. If/how the biofilm effects differ between the kaolin and MP?

## 2. The experiment

### 2.1 Test species and its feeding movements

*Daphnia magna* is planktonic microcrustacean, ecologically important, common, and easily cultured in the laboratory. It is also a standard species in ecological and ecotoxicological testing, including feeding inhibition assays (McWilliam and Baird 2002). Daphnids have two doubly-branched antennae (frequently half the length of the body or more) that are used for swimming and flattened leaf-like limbs inside the carapace (thoracic legs) that produce a current of water which carries food and oxygen to the mouth and gills. Small particles (generally <50 μm) are filtered out by fine setae on the thoracic legs and channeled along a groove at the base of the legs to the mouth. These animals feed on floating particles: phytoplankton, bacteria, and fungi, but also decaying organic material. Although there is some evidence of preferential feeding on certain types of algae, it is generally believed that all particles of suitable size are ingested without any selective mechanism. When rough material or tangled masses are introduced between the mandibles, they are removed by spines on the first legs and then kicked out of the carapace by the post-abdomen. We were intrigued by the possibility to detect movements related to the filtering activity and removal of the non-edible material (MP and kaolin) using electromagnetic signals at different frequencies.

### 2.2 Culture conditions

We used *Daphnia magna*, test strain *Klon 5* Federal Environment Agency, Berlin, Germany, cultured in M7 medium (artificial lake water; OECD, 2004; OECD, 2008). The animals were kept in groups of ∼25 individuals in 2 L containers and fed a mixture of the green algae *Pseudokirchneriella subcapitata* and *Scenedesmus subspicatus* three times a week. For the experiments, adult females with body length 2-3 mm were used.

### 2.3 Test particles

Kaolin (Sigma-Aldrich; size 4 ± 1 μm, density 2.6 g/cc), a clay mineral, with soft consistency and earthy texture, was used as a source of natural particles that daphnids would encounter even in the most pristine systems. As MP, we used fluorescent green microspheres (Cospheric, Santa Monica CA, USA; 1-5 μm, 1.4 g/cc).

Bovine Serum Albumin (BSA) was used for coating of MP and kaolin following the existing protocols (Taghon 1982). The particles incubated with BSA (2 mg/ml) were stored in the fridge for 48 h to ensure a complete coating. Using Zetasizer Nano Z (Malvern Instruments), zeta potential was measured in the non-coated and coated particles suspended in M7 at room temperature in folded capillary cell (DTS1060) at a modulator frequency of 1,000 Hz. The cell was rinsed with M7 medium between the samples to minimize contamination.

### 2.4 Exposure system and treatments

Multispecies Freshwater Biomonitor^®^ (MFB LIMCO; LimCo International, Germany) was used for movement recording. The MFB measures the activity of test animals by sending high frequency signals from one pair of electrode to another in a small cage. The cages were closed at both ends with nylon gauze (1 mm) fastened by screw-on rings (Figure 1A). Each cage contained a single *Daphnia* and was immersed in a separate 600-ml aquarium filled with M7 medium (Figure 1B). The exposures were conducted at room temperature and ambient light.

**Figure 1.**
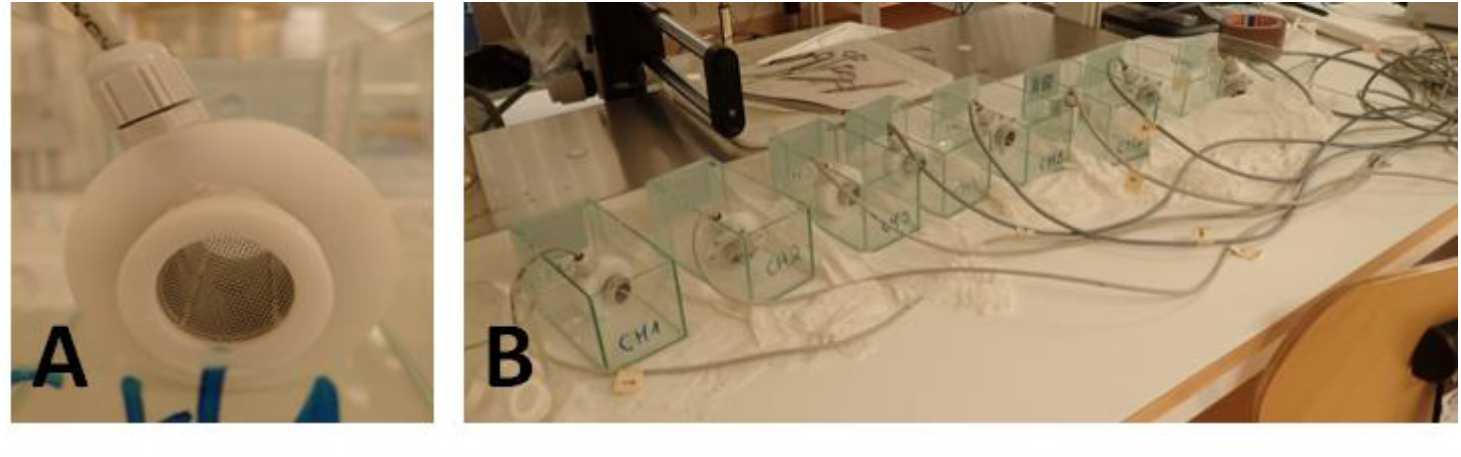
MFB test chamber for a single animal exposure (A) and the experimental aquaria used for simultaneous measurements with 8 channels (B). Quadropole impedance conversion technique based on the animal functioning as resistant in an alternating current between electrodes at opposite walls of the test chamber. Movements of the animal change the conductivity and the electrical field between a second, non-current carrying pair of electrodes and generate specific electrical signals for different kinds of behavior (swimming, locomotion, etc.).

The experimental design was 2 × 2 with particle type (Kaolin vs. MP) and coating (coated vs. non-coated) as treatment factors. The particle concentrations were 1.34 × 10^5^ and 1.45 × 10^5^ for non-coated and coated MP, respectively, and 1.12 × 10^5^ and 1.01 × 10^5^ for non-coated and coated kaolin, respectively. The concentrations of particles and algae were determined by Spectrex laser optical counter (SPECTREX Inc. USA). Test particles and algae (5 × 10^5^ cells/ml) were added to the aquaria at the start of the incubation. The eight experimental units were run simultaneously; 4 of them served as controls (only algae and daphnids) and 4 as particle tests (MP or kaolin, algae and daphnids). Algae and test particle settling at the bottom of the aquaria were resuspended on a few occasions during the exposure using a pipette. Upon termination of the exposure, the daphnids were retrieved and their body length was measured. No mortalities were recorded.

### 2.5 MFB readings and endpoints

*Daphnia* movements caused changes in the amplitude of the electrical signals generated at certain frequencies. The signal is depending on the type of activity and the size of the animal. In our pilot experiments, the MFB recordings were positive only in the range of 0.5-3.5 Hz; therefore, only this range was considered. A quantitative recording of the movements was made automatically for 12 h, starting every 10 min, with a trace of 10 sec. The raw data files containing the MFB records were analyzed according to the signal amplitudes and frequencies by discrete fast Fourier analysis (Gerhardt et al. 1998).

Based on the set of pilot experiments with various chemical substances known to affect specific motion types (caffeine, diclofenac, CO_2_-saturated water), we defined the filtering activity as corresponding to the routine low-amplitude movement of the thoracic legs and recorded at 0.5-1.0 Hz, whereas movements recorded at 2.0-3.5 Hz represented more chaotic and agitated movements related to high-amplitude strokes by antenna as well as discarding of non-edible material by post-abdominal claws and cleaning of the feeding appendages.

### 2.6 Data analysis and statistics

First, the recorded signals were averaged for 0.5-1.0 Hz (Filtering activity) and 2.0-3.5 Hz (Jumping activity). Then, the average proportions of time spent on Filtering activity and Jumping activity over the observation period were calculated using 10-sec observations over a 12 h period.

The values for Filtering and Jumping activities were Box-Cox transformed to approach normal distribution and normalized to the *Daphnia* body length. The resulting values were normalized to the respective values in the controls (*Daphnia* fed only algae and incubated during the same run). For statistical comparisons, we used two-way repeated measures ANOVA with either *Particle type* and *Coating* (when testing effects on zeta potential) or *Particle type* and S*wimming proxy* (when testing effects on swimming behavior) as fixed factors after the assumption of normality and homogeneity of data were met using Levene test. Individual-based subject matching was applied in the ANOVA analysis, because Filtering and Jumping activities are not independent within a time budget of an individual. Tukey’s test was used for pair-wise comparisons. Data are shown as means and standard deviations, α was set at 0.05.

## 3. Results and discussion

### 3.1 Effect of coating on zeta potential

Zeta potentials revealed that both test particles had negative net charges, and the values were within the range of values found for algal prey. In both particle types, coating resulted in the increased zeta potential; however, this change was significant for MP only as indicated by the significant interaction effect (F_1,8_ = 10.16; p = 0.013; Figure 2A). There were significant differences in zeta potential between the non-coated kaolin and MP (q_8_ = 8.167; p < 0.001), but not between the coated particles (q_8_ = 1.792; p > 0.05).

**Figure 2.**
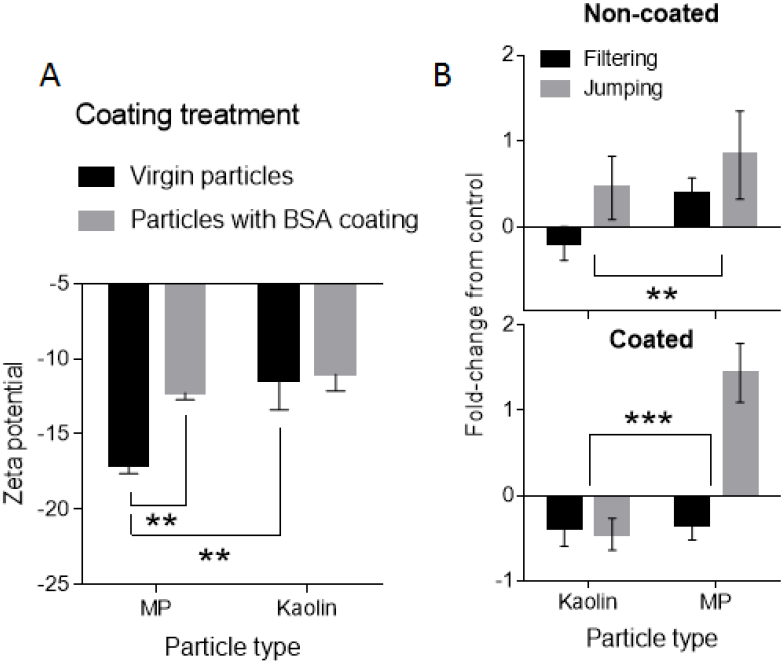
Experimental results for coating effects testing. (A) Effect of BSA coating on the surface charge measured as zeta potential (mean ± SD, n = 5) in kaolin and microplastics (MP). Tukey’s test was used for the pair-wise comparisons. (B) Swimming behavior of daphnids exposed to the test particles without assayed as Filtering and Jumping activities (mean ± SD, n = 4). The activity values show fold-change relative to controls (i.e., daphnids exposed to algal food only). Two-way RM-ANOVA comparison for *Particle type* is shown with asterisks; *: p<0.05, **: p<0.01; ***: p<0.001.

The coating was expected to result in a less negative charge and increased adhesive and aggregative capacities of both particle types (Taghon 1982, Matz and Jurgens 2001). This was indeed observed in MP, but not in kaoline; the latter was less negatively charged than MP, which might have been the reason behind the lack of its response to coating. However, given that both particle types with or without coating had weakly negative charge, they all were likely to be ingested by the animals. Nevertheless, significantly more negative charge in virgin MP implies that studies employing such particles may underestimate particle intake rate considering numerous reports on the relationships between particle retention rate and its surface charge in the various filter- and suspension-feeders (Taghon 1982).

### 3.2 Effect of test particles on swimming

The daphnids exposed to non-coated MP spent 25-35% more time for both Filtering and Jumping activities compared to those exposed to kaolin (Figure 2B, left panels). Although in the non-coated particle treatments none of the swimming proxies were significantly different from the controls, the difference between MP and kaolin was significant (F_1,6_ = 15.50, p = 0.008). In the coated particle treatments, Filtering and Jumping activities responded differently in the two particle types as indicated by a significant interaction effect (F_1,6_ = 84.11, p < 0.0001; Figure 2B, right panel). In the coated MP, the Jumping activity increased significantly compared to both coated kaolin (q_6_ = 13.31, p < 0.0001) and the controls (one-sided t-test, p < 0.05), whereas the Filtering activity was similar to that in kaolin and controls. Although the Filtering activity in KaolinBSA and MPBSA was not significantly different from the controls, the daphnids exposed to any test particle coated with BSA spent on average 35% less time filtering compared to those fed algae in the absence of the non-edible particles or with non-coated particles (Figure 2B).

These findings suggest that both kaolin and MP may alter swimming behavior yet in opposite way, with a slight decrease of filtration-related movements in kaolin and their increase in MP. Moreover, the presence of coating resembling biofilm was found to amplify this difference between the particles. The observed differences imply that daphnids would spend more time performing the non-feeding movements when swimming in suspension with MP, and even more when the MP are carrying biofilm. The increased jumping activity of filtrators exposed to microparticles (both natural and anthropogenic) may translate into changes in energy balance and growth. *In situ*, all particles would have much more developed biofilms compared to BSA coating produced in the lab. Moreover, these biofilms facilitate particle aggregation and thus increase the size of the particulates in the water, which could affect interactions with consumers and filtering efficiency (Rummel et al. 2017). Therefore, it is possible that stronger biofilm effects on the swimming behavior would be expected in weathered MP with heavy biofilm burden.

It is necessary to note that in this experiment we used very high concentrations of the test particles; such concentrations would correspond to highly turbid waters with a low ratio between the densities of algae and inert particles. At such densities, both natural mineral particles and synthetic polymers are likely to exert negative effects on various filter-feeder species by decreasing feeding efficiency (Boenigk and Novarino 2004). Nevertheless, the differences between responses to kaolin and MP indicate more severe swimming alterations and potentially higher foraging costs due to >2-fold increase in jumping and corresponding decrease in filtering activity brought about by presently unknown MP-animal interactions. The nature of these effects and the importance of particle physicochemical properties is still unclear and requires further investigation.

In previous studies on *Daphnia* movement analysis, swimming speed has been the favorite variable, but some other measures of swimming behavior have also been proposed (Dodson et al. 1997). Here, we report two useful proxies for daphnid swimming behavior related to food intake and time budget for feeding. Together they provide an efficient way for rapid screening of alterations in feeding behavior that are detectable long before the responses in food intake and growth can be detected.

Finally, separation of the animal’s foraging habits and behavior from its environmental setting is artificial, and the results can be misleading. While plastic litter is indeed a source of environmental pollution that should be combated, the immediate danger of microplastics for aquatic animals is often overstated. The effects reported so far are largely related to the effects that would be caused by any inert particles, such as those ubiquitously present in natural waters, and the downstream effects on food intake, growth, and reproduction. Therefore, intelligent testing of MP effects should include methods and techniques that help to understand the mechanisms behind these effects and delineate the responses to MP and those to naturally occurring suspended solids that occur in turbid waters.

